# Understanding the development of neural abnormalities in adolescents with mental health problems: a longitudinal study

**DOI:** 10.1101/2024.07.30.605786

**Authors:** Jiangyun Hou, Laurens van de Mortel, Weijian Liu, Shu Liu, Arne Popma, Dirk J.A. Smit, Guido van Wingen

## Abstract

Many mental health problems are neurodevelopmental in nature and have an onset during childhood. Mental health disorders are associated with neural abnormalities, but it is unclear when those emerge and how this relates to the development of different mental health problems. We used data from the largest longitudinal neurodevelopmental study to identify the structural and functional brain changes that co-occur with the onset of six mental health problems. The results showed premorbid brain-wide abnormalities that were comparable between internalizing and different from externalizing problems, and differential neurodevelopmental trajectories for specific brain regions in 11- to 12-year-old adolescents who developed ADHD, conduct, depressive and oppositional defiant problems. These results reveal that the onset of different mental health problems co-occur with common as well as problem-specific brain abnormalities.

## Introduction

Many mental health problems are neurodevelopmental in nature and have an onset during childhood or adolescence ^1,2^. Recent research suggests that approximately 22% of the general population is impacted by a mental disorder ^3^, and nearly one in seven adolescents meet the criteria for a psychiatric diagnosis ^4^. The prevalence of mental health issues among adolescents has even started rising in recent years ^5^, which makes it even more urgent to understand how mental disorders develop.

A common hypothesis states that mental disorders may develop during adolescence due to the marked neurodevelopmental changes ^6^. An increasing number of studies are employing longitudinal investigations to capture ongoing alterations and developments in the course of various disorders ^7–9^. Indeed, studies have shown altered neurodevelopmental trajectories in individuals with ADHD ^10,11^ and depression ^12^. But those studies are conducted in participants that have already developed a disorder. It therefore remains unclear whether those neurodevelopmental differences reflect a preexistent vulnerability or a consequence of having developed mental health problems. In order to disentangle cause from consequence, longitudinal studies are required to assess participants in the prodromal phase. Meanwhile, these studies often investigate single disorders, ignoring the fact that mental disorders often share symptoms and etiology. It therefore remains unknown whether differential neurodevelopmental trajectories could explain the vastly different clinical manifestations, or whether overlapping neuroanatomical changes are related to the development of mental health problems in general.

To investigate this, we selected participants from the Adolescent Brain Cognitive Development (ABCD) study ^13^, the largest longitudinal neuroimaging study focused in a critical developmental period from age 9-10. Our sample consisted of individuals who were initially healthy but later developed a range of mental health problems over a two-year follow-up period, and controls who remained healthy. Mental health problems were identified based on DSM-5 criteria and categorized into six distinct types: ADHD, anxiety, conduct, depressive, oppositional defiant, and somatic problems as assessed with the Child Behavior Checklist (CBCL 6-18) ^14^. We extracted structural and functional neuroimaging data collected at baseline and at two-year follow-up and investigated whether neural abnormalities already existed at baseline or developed together with the onset of mental health problems.

## Results

We compared six groups that consisted of participants who developed different mental health problem (ADHD, anxiety, conduct, depressive, oppositional defiance, and somatic problems; N=58-173) with controls (N=2500) without clinically relevant symptoms. There were several significant differences in demographic data between groups (Table 1). To account for differences in demographic composition, all analyses were corrected for age, sex, IQ and parental EA. In addition, we performed sensitivity analyses in which we matched the control group on these characteristics, for which the results remained comparable (see Supplementary Fig. S3-S6, Supplementary Table S5).

**Table 1.**
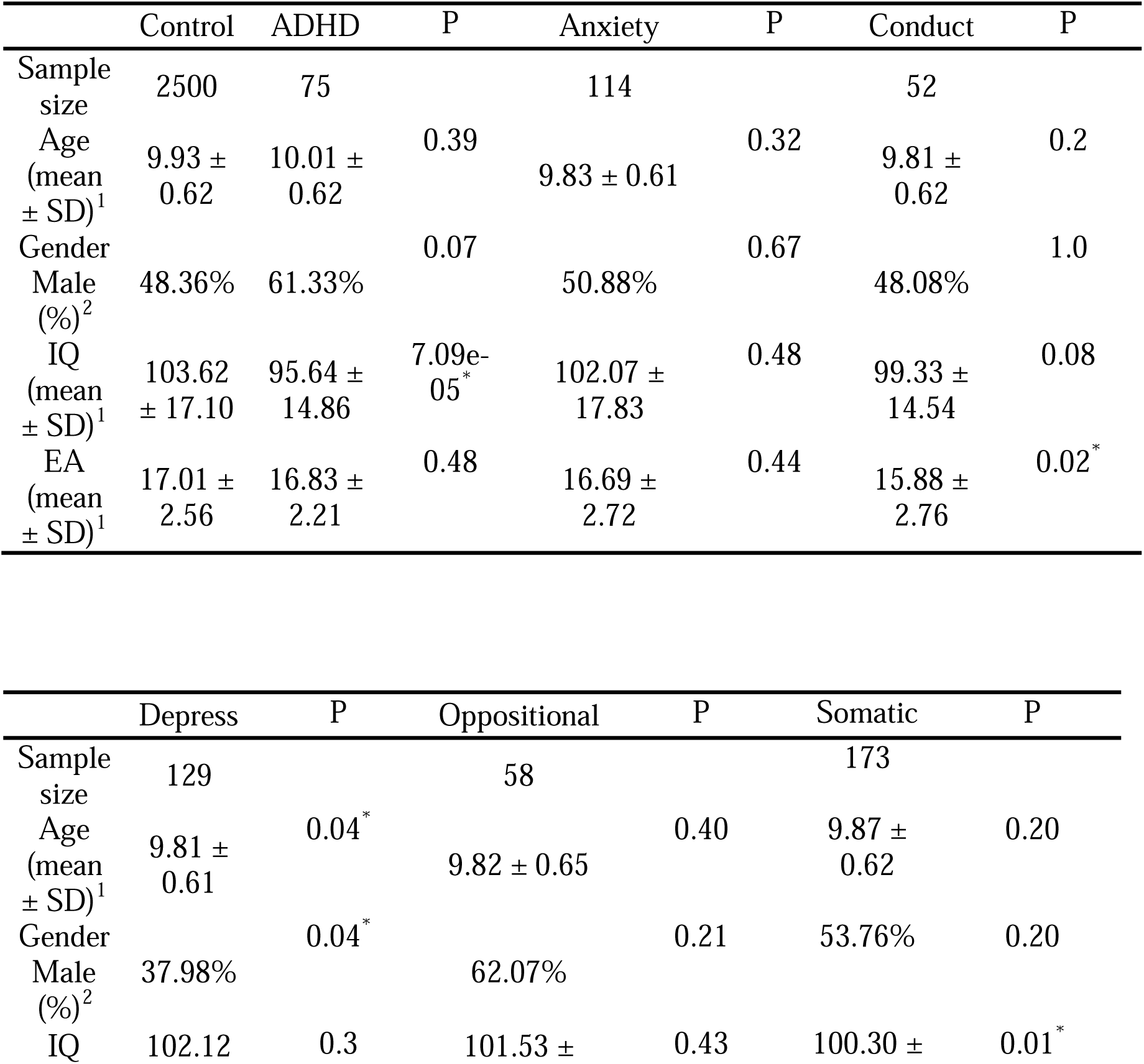
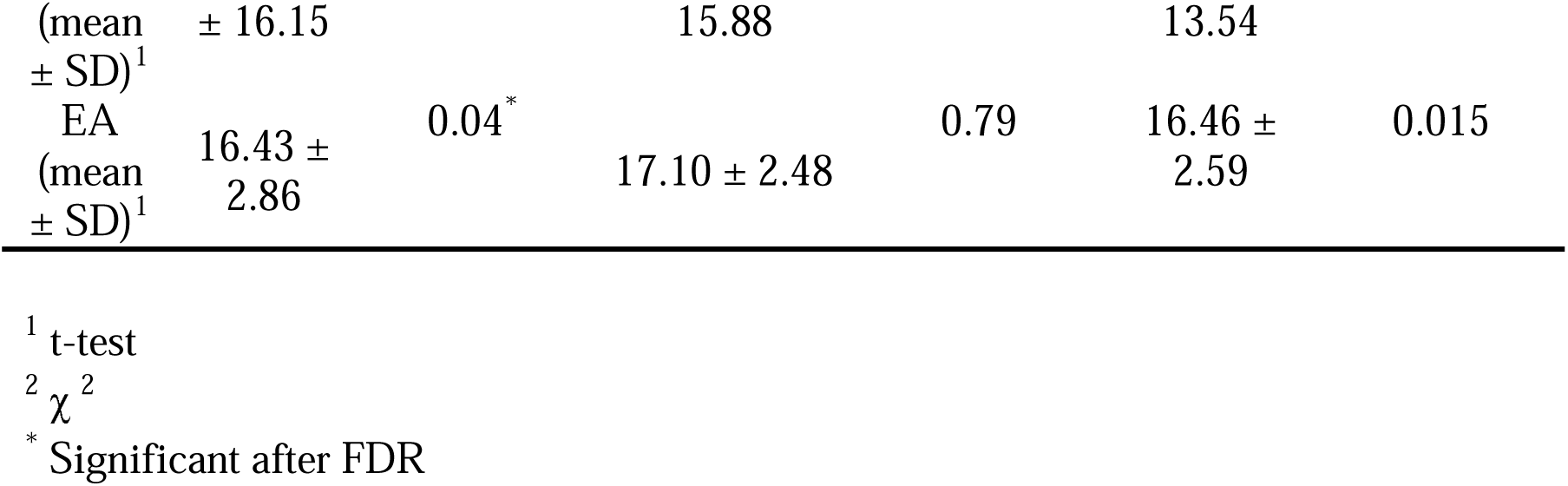
Demographic data of the seven adolescents’ groups at baseline. The six disorder groups included participants without clinically relevant symptoms at baseline (CBCL t-score < 65) but with clinically relevant symptoms at two year follow-up (CBCL t-score ≥ 65). Controls did not have clinically relevant symptoms at baseline nor at follow-up (CBCL t-score < 65). P values are false discovery rate corrected.

We used imaging data from six modalities at baseline and two year follow up to investigate neurodevelopment from no clinical symptoms to clinically relevant mental health symptoms: sMRI, rsfMRI, DTI, and task-based fMRI. Experimental paradigms included the monetary incentive delay (MID) task to assess reward processing, the stop signal task (SST) to assess behavioral inhibition, and the emotional n-back (N-back) task to assess emotional working memory.

### Baseline

After false discovery rate (FDR) correction for multiple comparisons, we observed significant differences at baseline (before the onset of clinically relevant symptoms) for the conduct and oppositional defiant problem groups compared to healthy controls in rsfMRI measures (**Fig. 1**., Supplement Fig.S1), but not for ADHD, anxiety, depressive and somatic problem groups. The conduct problem group showed increased temporal variance in the left superior frontal gyrus at rest. The oppositional defiant problem group showed increased temporal variance in the left superior temporal gyrus, left superior frontal gyrus and left lateral occipital gyrus at rest (Supplement Table S2). We found no significant differences at baseline for any of the other neuroimaging measures. Thus, only individuals that later developed conduct or oppositional defiant symptoms already had prodromal regionally specific differences in rsfMRI measures.

**Fig. 1.**
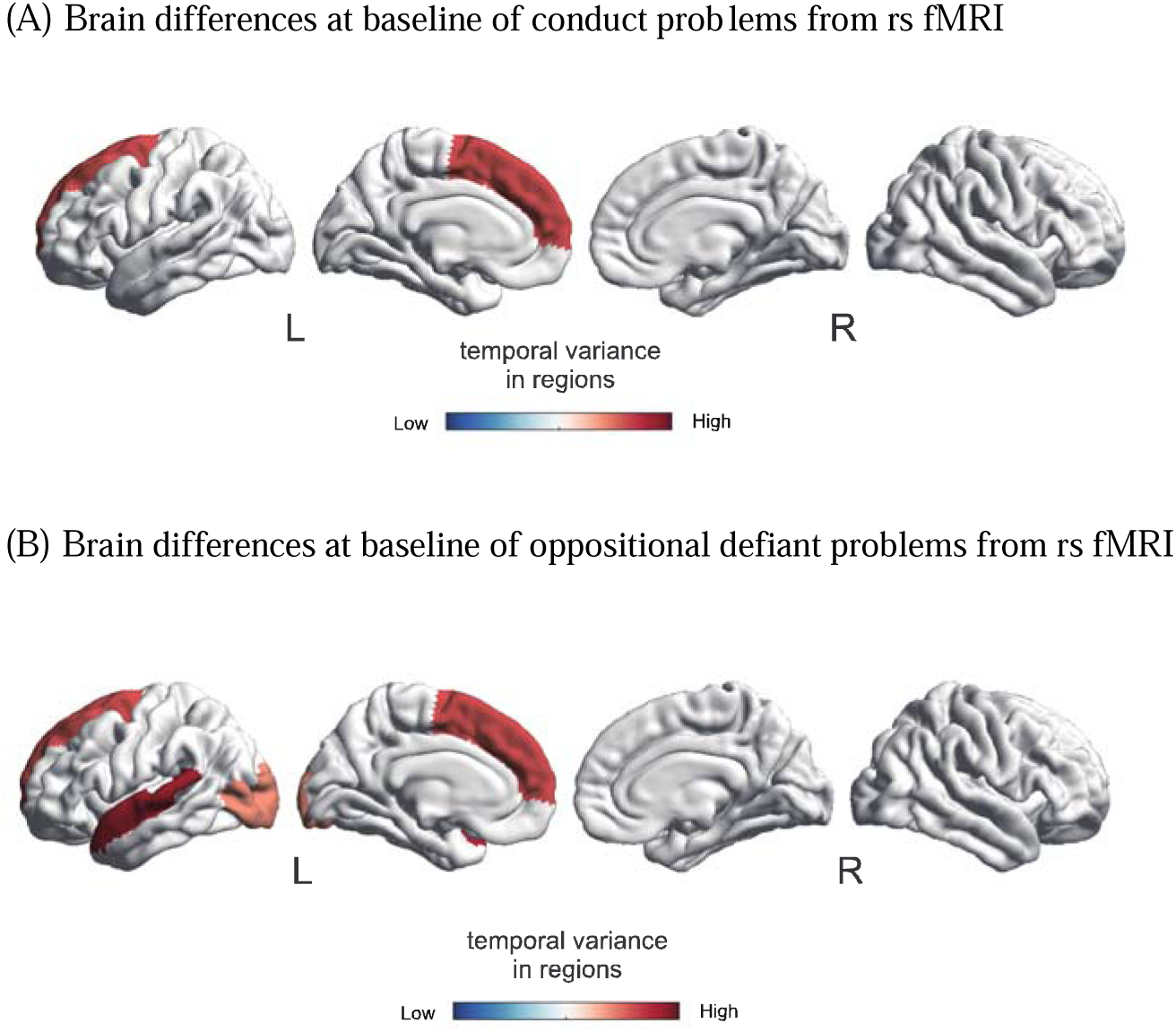
Aberrant brain regions in conduct and oppositional defiant problems at baseline. (A) Conduct: increases in cortical rsfMRI variance, (B) Oppositional defiant: increases in cortical rsfMRI variance.

### Developmental trajectories

We next tested whether there were significant differences in neurodevelopment between the different symptom groups and healthy controls. We found differential neurodevelopmental changes at two-year follow-up (after the onset of clinically relevant symptoms) for ADHD, depressive, conduct, and oppositional defiant problem groups (**Fig. 2**.; Supplement Fig.S2). For ADHD problems, the results in this group showed higher resting-state functional connectivity between the ventral attention network (VAN) and right caudate nucleus. The depressive problem group showed increased temporal variance in the right thalamus. For conduct problems, rsfMRI measures showed increased temporal variance in the right middle temporal gyrus, left posterior cingulate cortex, right lingual gyrus, left supramarginal gyrus, left fusiform gyrus, left lingual gyrus, and right superior temporal gyrus. In addition, this group showed decreased activity during the SST for correct stop versus incorrect stop in the left pallidum and right ventral diencephalon, and for correct stop versus correct go contrast in the left pallidum. In the oppositional defiant problem group, results showed increased left cerebral white matter volume, and increased fiber tract volume of all white matter tracts combined, and the majority of the individual fiber tracts (see Supplement Table S3). While no statistically significant differences were observed for the anxiety and somatic problem groups, some of the features showed a trend towards statistical significance (Supplement Fig. S2, Supplement Table S4). These results indicate that the ADHD, conduct, oppositional defiant, and depressive problem groups had different neurodevelopmental changes after the onset of mental health problems, while no significant effects were observed for the anxiety and somatic groups.

**Fig. 2.**
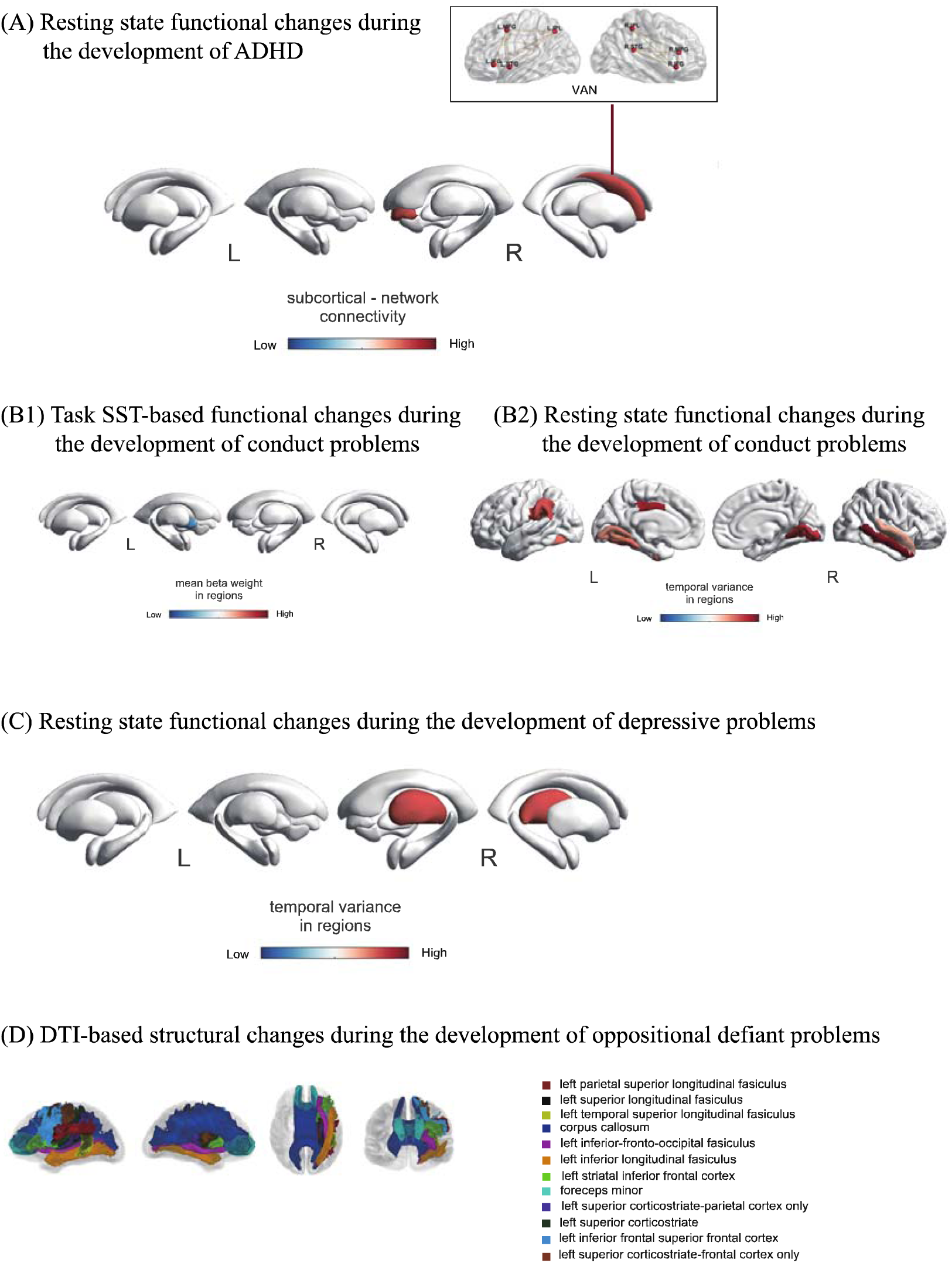
Aberrant neurodevelopment in ADHD, conduct, depressive, and oppositional defiant problems. (A) ADHD: increases in subcortical functional connectivity; (B1) Conduct: decreases in subcortical regions during performance of the SST, (B2) increases in cortical rsfMRI variance; (C) Depression: increases in thalamic rsfMRI variance; (D) Oppositional defiant: increases in DTI tract volume.

### Correlations

To evaluate whether the neurodevelopmental changes were shared between symptom groups or specific per group, we performed pairwise brain-wide correlations for each MRI modality separately between the different mental health problem groups using t-values across the brain for the comparison with controls ^15^. At baseline and after FDR correction for multiple comparisons, the results showed opposing patterns of brain abnormalities before the onset of clinically relevant symptoms (**Fig. 3**.). While the ADHD, depressive, anxiety and somatic problem groups largely showed positively correlated brain differences compared to controls, the conduct and oppositional defiant problem groups showed negatively correlated brain differences with the other groups. Specifically, the oppositional defiant problem group showed negative correlations with several other groups in DTI, sMRI and task-related activity during the MID and emotional N-back task, and the conduct problem group showed negative correlations with other groups during the MID. Interestingly, the oppositional defiant and conduct problem groups also showed a negative correlation for sMRI, indicating that structural brain abnormalities were specific to the oppositional defiant problem group.

**Fig. 3.**
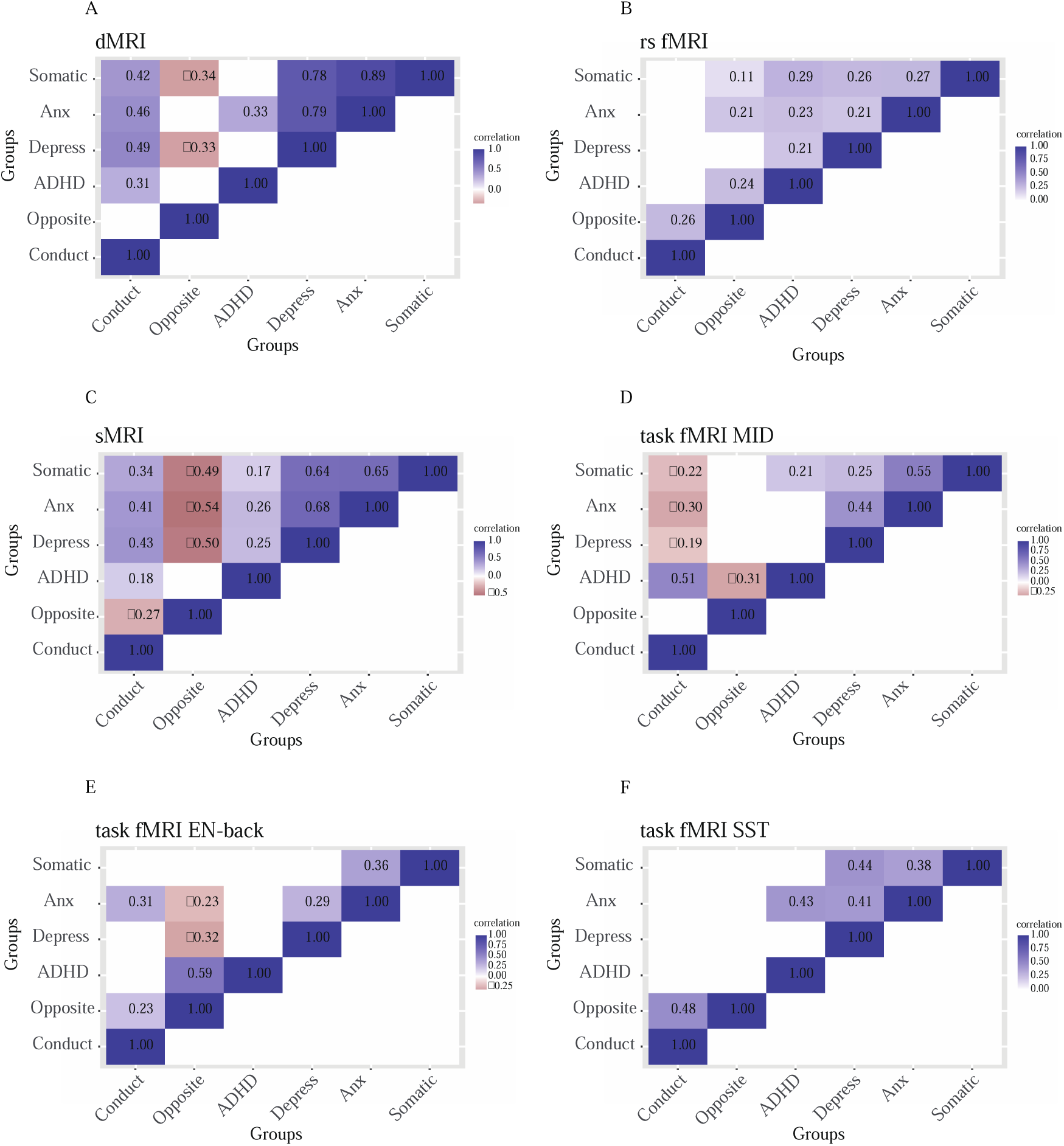
Correlation matrices between mental health problems in different modalities at baseline (blue is positive and red is negative). Only significant correlations between t-values from the baseline linear models at p(FDR)<0.05/6 are shown. (A) DTI; (B) rsfMRI; (C) sMRI; (D) MID task fMRI; (E) EN-back task fMRI; (F) SST task fMRI.

We performed the same correlation analysis for the changes in MRI modalities at two-year follow-up (after the onset of clinically relevant symptoms), which largely showed positive correlations for each of the group pairs and MRI modalities (**Fig. 4**.). The majority of correlations between groups were significant for rsfMRI and structural MRI, while approximately half the correlations were significant for DTI and task-based fMRI. Notable exceptions were negative correlations in DTI changes between ADHD and depressive problem groups, and activity during the MID task between the anxiety and oppositional defiant problem groups.

**Fig. 4.**
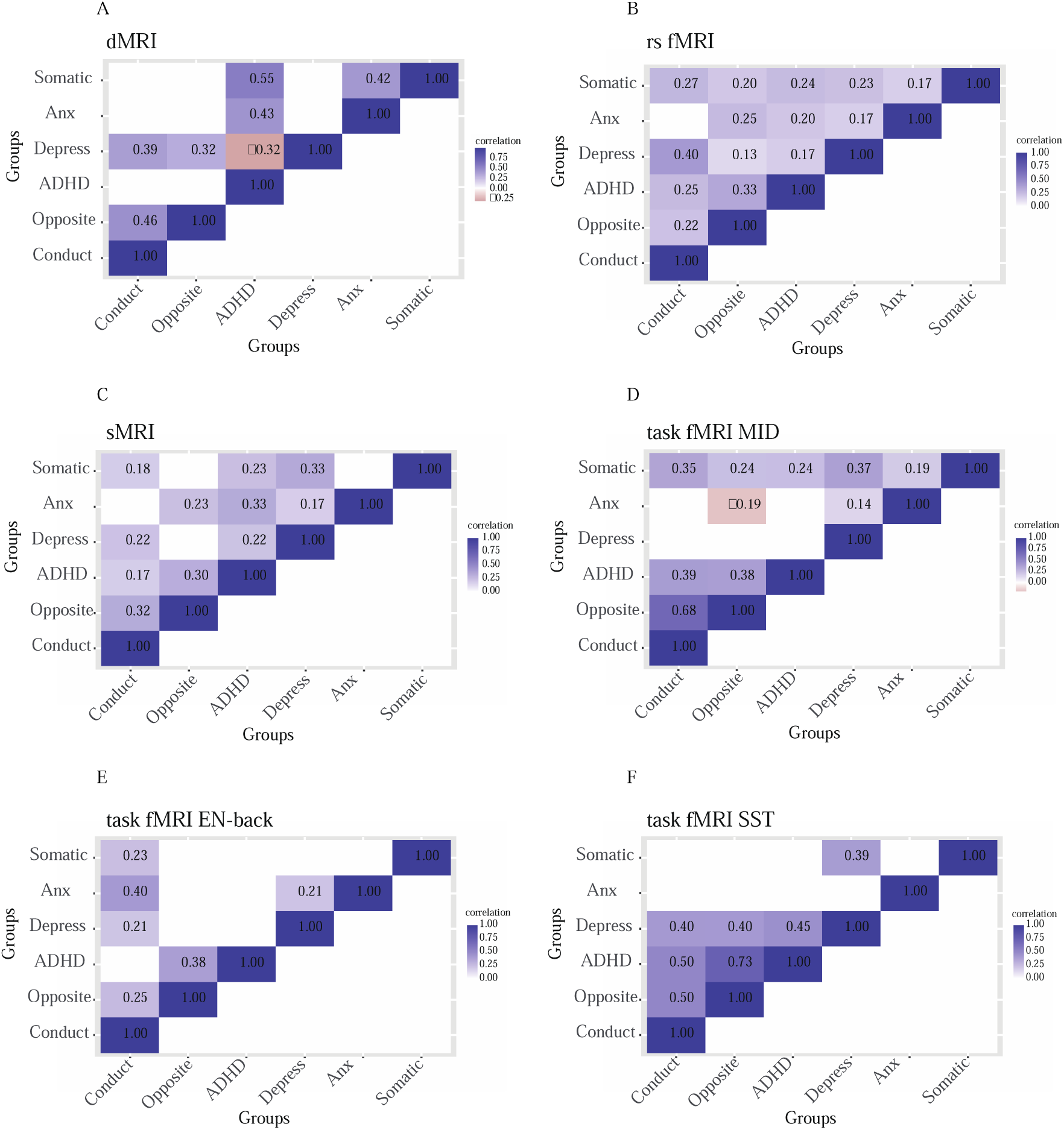
The correlation matrices between mental health problems in different modalities of changes (blue is positive and red is negative). Only significant correlations between t-values from the change linear models at p(FDR)<0.05/6 are shown. (A) DTI; (B) rsfMRI; (C) sMRI; (D) MID task fMRI; (E) EN-back task fMRI; (F) SST task fMRI.

## Discussion

This study investigated whether neural abnormalities already exist before the development of different mental health problems or whether those develop with the onset of mental health problems. Our results show that the onset of mental health problems in youth between 9 to 12 years of age is primarily associated with altered neurodevelopment, with limited prodromal differences in brain structure and function. The longitudinal comparison to controls revealed neurodevelopmental changes that were specific for different mental health problems, while the correlation analysis suggests that most of altered neurodevelopment was shared between symptom groups. Together, these results suggest that structural and functional brain abnormalities primarily emerge in parallel during the development of clinically relevant mental health problems, and that these contain both common altered brain-wide neurodevelopmental trajectories as well as brain changes that are specific to particular symptoms.

Participants with ADHD, conduct, depressive and oppositional defiant problems showed significant differences in neurodevelopment compared to healthy controls. ADHD, depressive, and conduct problems were associated with differential changes in resting-state fMRI. The conduct problem group also showed differences in activity during behavioral inhibition, while oppositional defiant problems were associated with additional wide-spread differences in white matter. Thus, neural differences in white matter and during behavioral inhibition were specific for conduct and oppositional defiant symptoms. While the changes in resting-state functional connectivity across the brain were largely comparable between groups, the results also showed specific changes. Problem-specific changes were limited to the subcortex (thalamus) for depressive problems, to the cortex for conduct problems limited, and to cortical-subcortical interactions for ADHD problems. No significant differences were observed in individuals with anxiety and somatic problems, which may only evolve after longer disease duration.

The neuroimaging results for the specific groups are in line with what has been previously reported for patients with longer disease durations, for whom it is unclear whether neural abnormalities were already present during the premorbid phase, related to the onset of symptoms, or a consequence of having a disorder (scarring). In this study, at baseline, we found increased temporal variance of the left superior frontal gyrus in the conduct problem group. Alterations of the superior frontal gyrus in both structure and function have been reported previously in conduct problems^16–20^. The left superior frontal gyrus was also found to have increased temporal variance at baseline in the oppositional defiant problem group, along with the left superior temporal gyrus and left lateral occipital gyrus, which could reflect commonalities in functional alterations between the conduct and oppositional defiant problem groups. The left superior temporal gyrus and lateral occipital gyrus have shown alterations in individuals with combined ADHD and oppositional defiant problems^21,22^, although the results from these studies were based on individuals which already developed conduct or oppositional defiant problems. Our results indicate that these cortical changes occur before the onset of many symptoms, which may help identify early signs of conduct and/or oppositional defiant problems. In addition, these significant differences were found only in externalizing problems at baseline, which may indicate that the functional neural abnormalities in externalizing problems occur earlier than in internalizing problems.

During the development of problems, we found higher functional connectivity between the VAN and the caudate nucleus in ADHD problem group. The VAN, which consists of regions within the ventral prefrontal cortex and the temporoparietal junction, can redirect attention toward salient stimuli, particularly novel stimuli ^23^. Hypofunction of the VAN has previously been reported in ADHD ^24^. The caudate nucleus plays an important role in motor, emotional, and cognitive functions ^25^, and the dysfunction of it has been associated with ADHD ^26^. Although no study has directly reported altered connectivity in adolescents with ADHD problems between the VAN and caudate nucleus, this altered connectivity may underlie the attentional and executive dysfunctions which are characteristic of ADHD. During the development of conduct problems, we found increased temporal variance in seven cortical regions. These regions have previously shown differences in brain structure and function in individuals with conduct problems^27,28,29,30,31,32^. In addition, we observed decreased activity in the pallidum in this group during behavioral inhibition. The pallidum is involved in the regulation of voluntary movement, inhibitory control and various cognitive and emotional functions ^33–35^ and decreased pallidum activity during behavioral inhibition might be related to the impulse control impairment associated with conduct problems.

In the depressive problem group, we found increased brain activity in the right thalamus at rest. Previous studies have highlighted the role of the thalamus in depression. It is the most important brain region for the identification of pediatric and adult depression, ^36,37^ which is involved in emotional salience regulation ^38^. Our findings suggest an important role for the thalamus in the development of depressive problems, which may reflect increased processing demands for emotional salience regulation during the onset of depressive symptoms. Finally, we also found increased cerebral white matter volume during the development of oppositional defiant problems, which is consistent with a previous study ^21^. However, differences in fiber tract volume have rarely been reported for oppositional defiant problems, and the observed differences in fiber tract volume may be a new direction for studying the neurodevelopmental trajectory of oppositional defiant problems at an early stage. Our results indicate that the above neural abnormalities develop during the onset of clinically relevant symptoms, rather than that those reflect premorbid abnormalities or scarring effects.

While the longitudinal comparison to healthy controls shows that multiple neural abnormalities are specific to particular mental health problems, the correlation analysis between symptom groups primarily showed positive correlations for all MRI modalities. This suggests that the different symptom groups also share an abnormal brain-wide neurodevelopmental trajectory. This concept fits well with the general structure of mental health problems, for which the most parsimonious model contains a general psychopathology factor with additional specific symptom clusters ^39^. Furthermore, the negative correlation between anxiety and oppositional defiant problem groups during the MID suggest opposite activity patterns between these two groups, and the negative correlation between ADHD and depressive problem groups for DTI suggest opposite neural structural changes between these groups. Together, our findings imply that altered brain development consists of a largely shared neurodevelopmental factor with additional symptom-specific brain changes.

While the comparisons between symptom groups and healthy controls hardly showed significant differences at baseline (before the onset of clinically relevant symptoms), the correlations between symptom groups at baseline did reveal abundant whole-brain correlation patterns of neural aberrancies. We found high correlations for structural (0.64-0.89) and moderate correlations for functional (0.21-0.55) abnormalities between groups with internalizing problems (depressive, anxiety and somatic problems) ^40^, which were considerably stronger than for the positive correlations after the onset of clinically relevant problems (0.14-0.42). This may suggest that there is considerable overlap in neuroanatomy between internalizing groups in the prodromal phase. But as these problems developed in different directions, the correlations between the neuroanatomical differences weaken. Strikingly, even negative correlations were observed for the two groups with externalizing problems. The DTI measures of the conduct problem group were negatively correlated with those for internalizing problems. And DTI, structural MRI, and brain activity measures during emotional working memory of the oppositional defiant group showed negative correlations with those for internalizing problems. This suggests that functional and structural brain abnormalities of the (externalizing) conduct and oppositional defiant groups already existed at baseline. Meanwhile, the findings for the internalizing groups reflect the comorbid nature of mental health disorders, as previous studies have shown that depression, anxiety problems and somatic problems often co-occur, sometimes with ADHD ^41–45^.

The current study is based on the large ABCD cohort study, which provides sufficient imaging data at baseline and follow-up for multiple mental health problems. Nevertheless, there are several limitations to this study. First, this study is based on children from North America, and whether the results also apply to children from other continents requires further investigation. Second, the children in this study were 9 to 10 years old at baseline, and we do not yet know whether the results also apply to children of younger or older age. Finally, we included individuals who also had other problems as the resulting sample was not sufficiently large to exclude participants with comorbid problems. While this is beneficial for the generalization of the results to the clinical population, comorbid symptoms may have limited the identification of differences between problem groups.

In conclusion, our longitudinal study reveals complex neurodevelopmental changes in adolescents who develop clinically relevant mental health problems between 9 to 12 years of age. They have premorbid neural abnormalities that are different for internalizing and externalizing problems, have altered brain-wide neurodevelopmental trajectories that are in common, and additional symptom specific brain changes. These results contribute to our understanding how neural abnormalities evolve and relate to the development of distinct mental health problems.

## Method

### Participants

The Adolescent Brain Cognitive Development (ABCD) Study, including around 11,876 young people from age 9 to 10 at baseline, aims to understand the neurodevelopmental trajectories in adolescence and their relations to behavior, environment, and genetics. By recruiting participants from various schools and communities across the United States, the study gathers a diverse cohort representing a broad spectrum of socio-economic, cultural, and geographic backgrounds. In the ABCD study, participants undergo comprehensive assessments, which include neuropsychological tests, brain imaging, and various questionnaires about behavior, health, and lifestyle. One of the key components of this battery to assess mental health problems is the Child Behavior Checklist (CBCL), providing critical information on participants’ emotional and behavioral functioning ^13^.

The CBCL is a widely used tool for assessing emotional and behavioral problems in children. The CBCL is designed to obtain a comprehensive picture of a child’s functioning from the perspective of their parent or primary caregiver. The questionnaire covers various domains, including social interactions, academic performance, emotional regulation, and behavioral tendencies. The CBCL scores can be categorized into different syndromes, which provide valuable insights into potential clinical conditions or developmental issues. The use of standardized scores allows for comparisons across children and helps clinicians and researchers pinpoint specific areas of concern.

We selected ABCD study participants with initial t-scores under 65 on the DSM-oriented CBCL scales, but who exceeded this threshold at two-year follow-up. This lenient criterion of 65 offers a balanced sensitivity-specificity trade-off ^46^. We identified 238 individuals with ADHD problems, 311 with anxiety problems, 181 with conduct disorder problems, 361 with depressive problems, 219 with oppositional defiant problems, 412 with somatic problems, and 4842 healthy controls (HC) with a t-score <65 at baseline and at one and two year follow-up. After excluding those with a psychiatric history at baseline (defined as having ever received mental health or substance abuse services) or without usable MRI data at baseline and/or follow-up (as large missing MRI datasets cannot be imputed), our final sample consisted of 74 individuals with first onset ADHD, 114 with anxiety problem, 52 with conduct problem, 129 with depressive problem, 58 with oppositional defiant problem, 173 with somatic problem, and 2500 controls (**Table1, Fig. 5**.). Data can be obtained via https://nda.nih.gov/abcd.

**Fig. 5.**
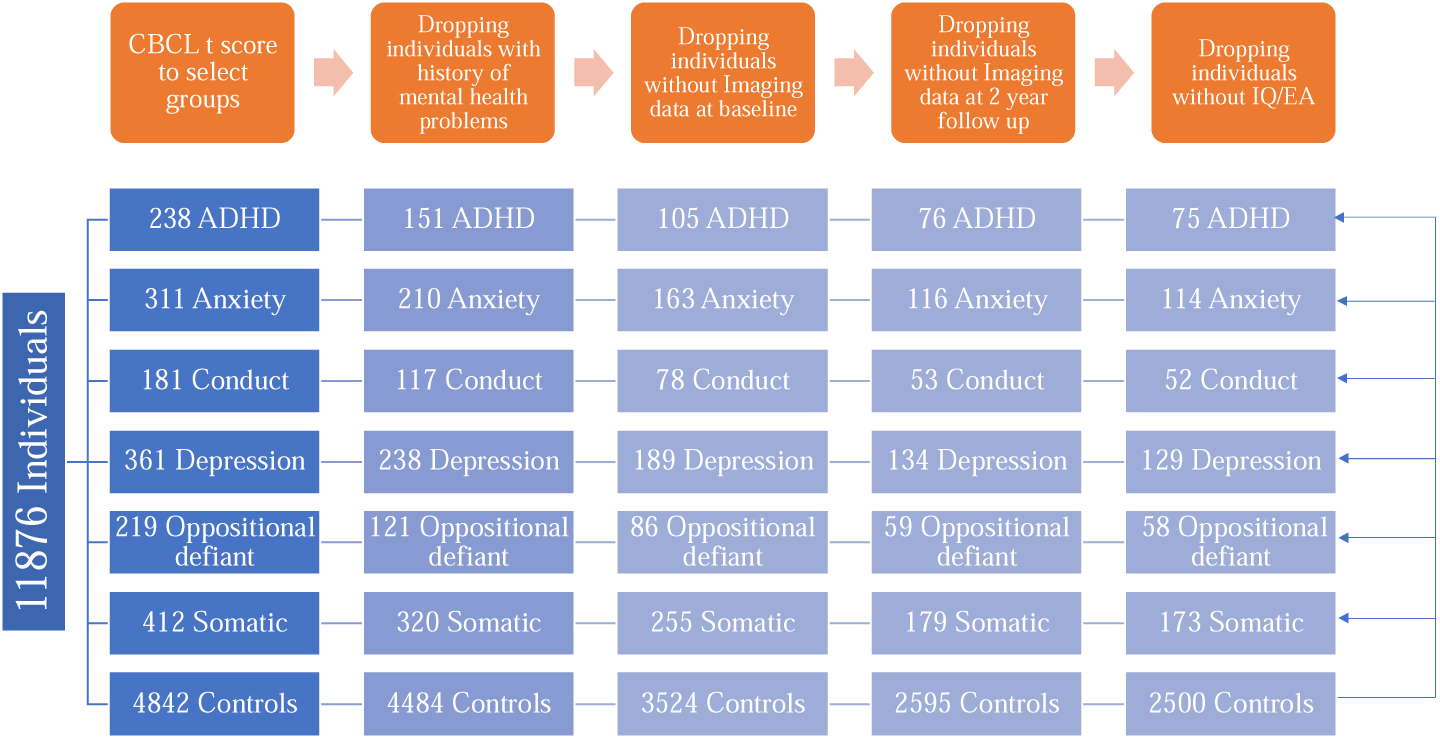
Flowchart illustrating the selection of research participants.

### Imaging data

Imaging data included 1876 image features from six MRI modalities extracted from the ABCD dataset using the published protocol: structural MRI, diffusion MRI, and functional MRI data (rsfMRI, task fMRI: MID, task fMRI: SST, task fMRI: EN-back)^47^. We selected 258 sMRI measures, 84 DTI measures, and 856 rsfMRI measures, 90 SST measures, 196 N-back measures, and 392 MID measures (Supplement Table S1). All preprocessing and analysis for these MRI measures were reported by Hagler et al ^47^.

### Analysis

Statistical analyses were performed in R. Separate linear models were used to assess group differences at baseline and differences in neurodevelopment for six disorders and six modalities between individuals with mental health problems and healthy controls with age, sex, IQ and educational level of parents (EA) as covariates. For baseline: baseline ∼ Group + Age +Sex +IQ + EA. For neurodevelopmental changes: two year follow up ∼ Group + baseline + Age +Sex +IQ +EA ^48^.

Statistical tests were false discovery rate (FDR) corrected for multiple comparisons across measures per imaging modality and Bonferroni corrected for testing multiple modalities for each disorder separately (p(FDR)<0.05/6). In addition, we assessed the degree of shared neurodevelopmental changes between mental health problems by estimating the correlation of t values across the brain from the baseline and change models for each neuroimaging modality separately (p(FDR)<0.05/6).

## Acknowledgments

This project was supported by a Chinese Scholarship Council scholarship awarded to YH. Data used in the preparation of this article were obtained from the Adolescent Brain Cognitive Development (ABCD) Study (https://abcdstudy.org), held in the NIMH Data Archive (NDA). This is a multisite, longitudinal study designed to recruit more than 10,000 children age 9-10 and follow them over 10 years into early adulthood. The ABCD Study is supported by the National Institutes of Health and additional federal partners under award numbers U01DA041048, U01DA050989, U01DA051016, U01DA041022, U01DA051018, U01DA051037, U01DA050987, U01DA041174, U01DA041106, U01DA041117, U01DA041028, U01DA041134, U01DA050988, U01DA051039, U01DA041156, U01DA041025, U01DA041120, U01DA051038, U01DA041148, U01DA041093, U01DA041089, U24DA041123, U24DA041147. A full list of supporters is available at https://abcdstudy.org/federal-partners.html. A listing of participating sites and a complete listing of the study investigators can be found at https://abcdstudy.org/consortium_members/. ABCD consortium investigators designed and implemented the study and/or provided data but did not necessarily participate in the analysis or writing of this report. This manuscript reflects the views of the authors and may not reflect the opinions or views of the NIH or ABCD consortium investigators.

## Authors contributions

Jiangyun Hou performed Methodology, Software, Validation, Visualization and Writing – original draft;

Laurens van de Mortel performed Methodology, and Writing – review & editing;

Weijian Liu performed Visualization;

Shu Liu performed Visualization;

Arne Popma performed Methodology;

Dirk Smit performed Methodology, and Writing – review & editing;

Guido van Wingen performed Methodology, Supervision, Writing – review & editing

## Data and materials availability

Data used in the preparation of this article can be obtained from the Adolescent Brain Cognitive Development (ABCD) Study (https://abcdstudy.org), held in the NIMH Data Archive (NDA).

