## Supplementary material for "Understanding the development of neural abnormalities in adolescents with mental health problems: a longitudinal study": Figs. S1to S6 Tables. S1 to S5

**Understanding the Development of Mental Disorders Through Longitudinal Neuroanatomy in Adolescents**

Guido van Wingen

Corresponding author: Jiangyun Hou and Guido van Wingen

**The file includes:**

Figs. S1to S6

Tables. S1 to S5

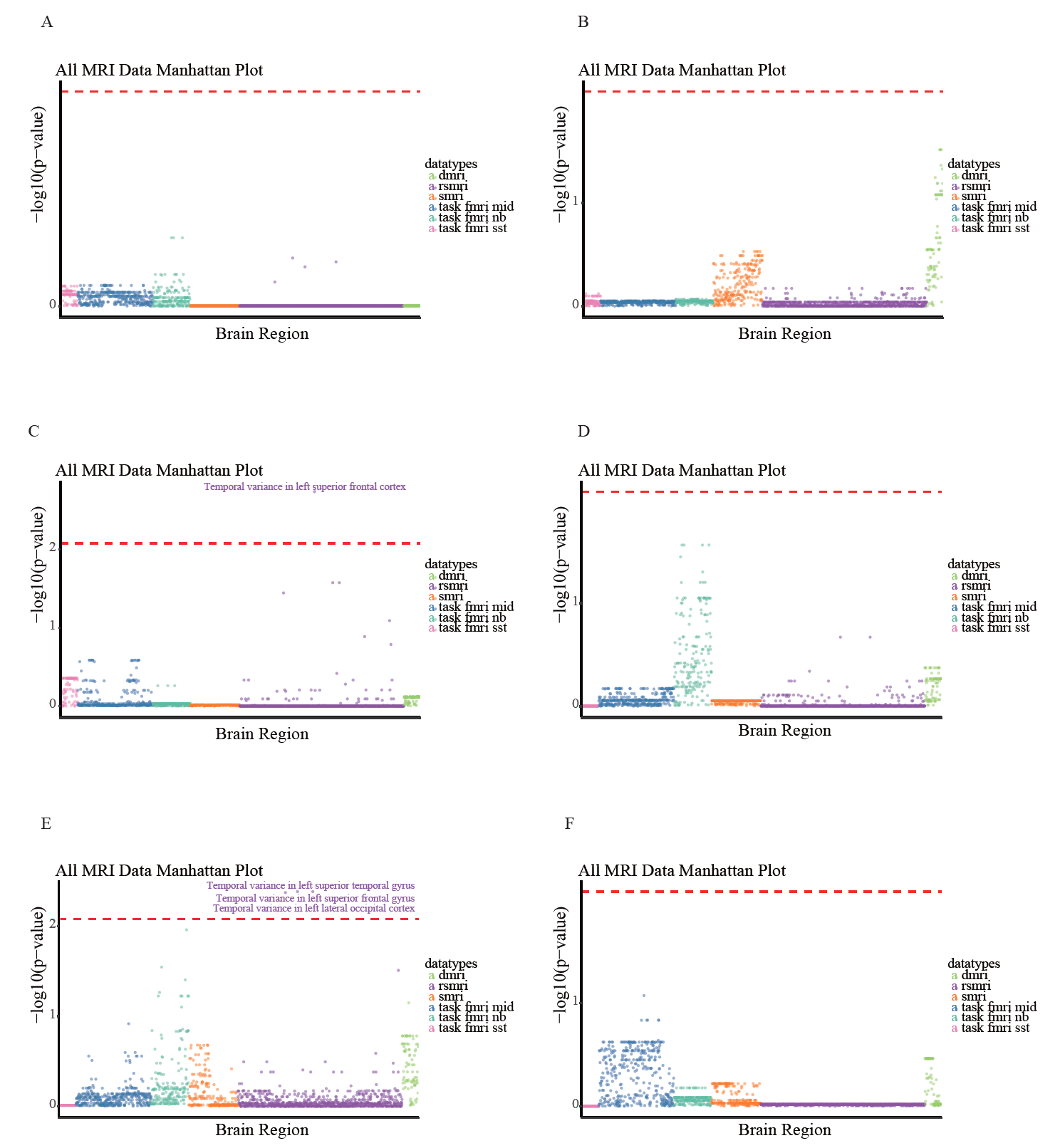

**Fig. S1. The results of linear model at baseline**. (A) the Manhattan plot of the baseline linear model results of ADHD with p(FDR)<0.05/6 (red line); (B) the Manhattan plot of the baseline linear model results of anxiety with p(FDR)<0.05/6 (red line); (C) the Manhattan plot of the baseline linear model results of conduct with p(FDR)<0.05/6 (red line); (D) the Manhattan plot of the baseline linear model results of depression with p(FDR)<0.05/6 (red line); (E) the Manhattan plot of the baseline linear model results of oppositional defiant with p(FDR)<0.05/6 (red line); (F) the Manhattan plot of the baseline linear model results of somatic with p(FDR)<0.05/6 (red line).

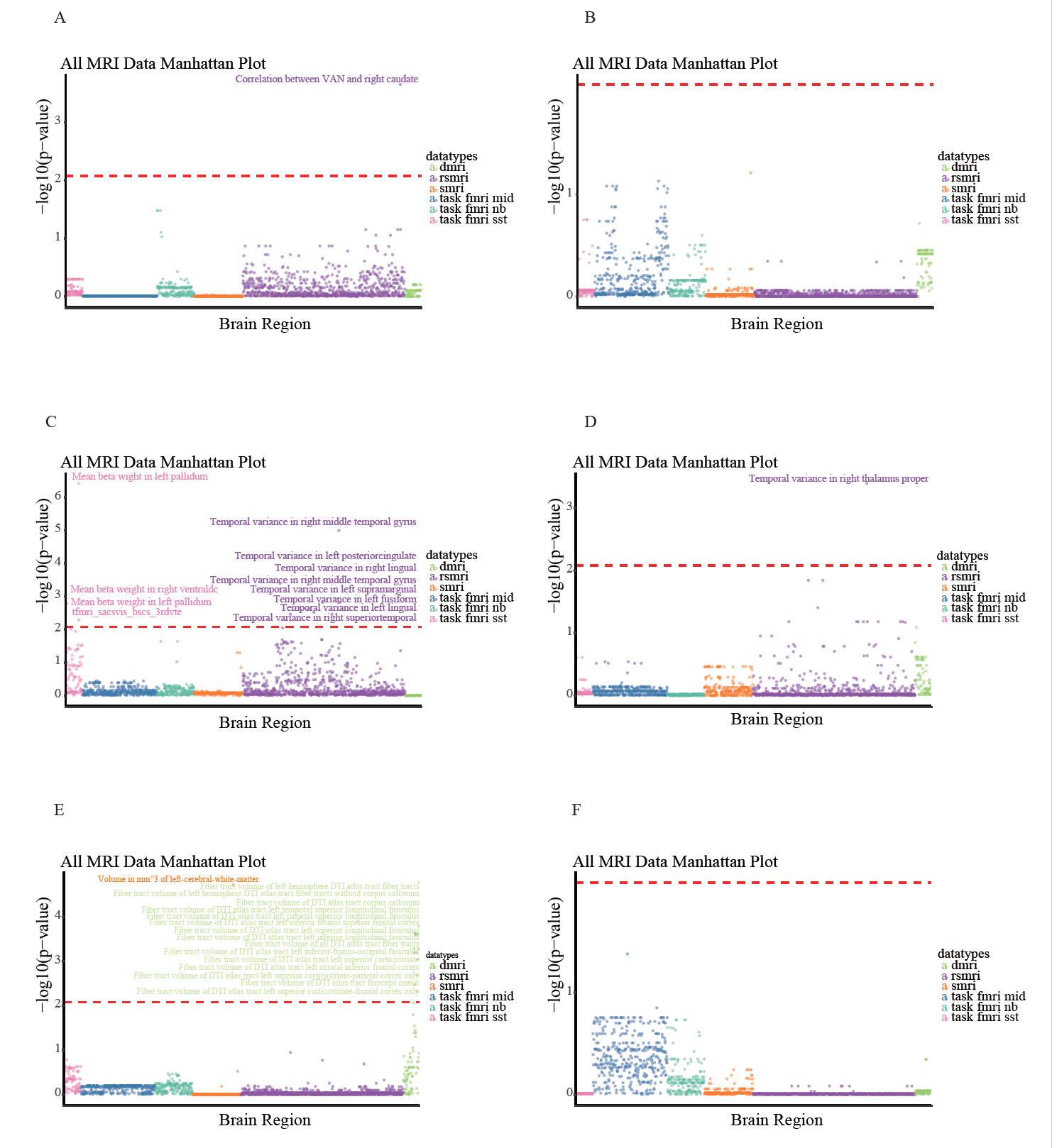

**Fig. S2. The results of linear model of changes**. (A) the Manhattan plot of the change model results of ADHD with p(FDR)<0.05/6 (red line); (B) the Manhattan plot of the change model results of anxiety with p(FDR)<0.05/6 (red line); (C) the Manhattan plot of the change model results of conduct with p(FDR)<0.05/6 (red line); (D) the Manhattan plot of the change model results of depression with p(FDR)<0.05/6 (red line); (E) the Manhattan plot of the change model results of oppositional defiant with p(FDR)<0.05/6 (red line); (F) the Manhattan plot of the change model results of somatic with p(FDR)<0.05/6 (red line).

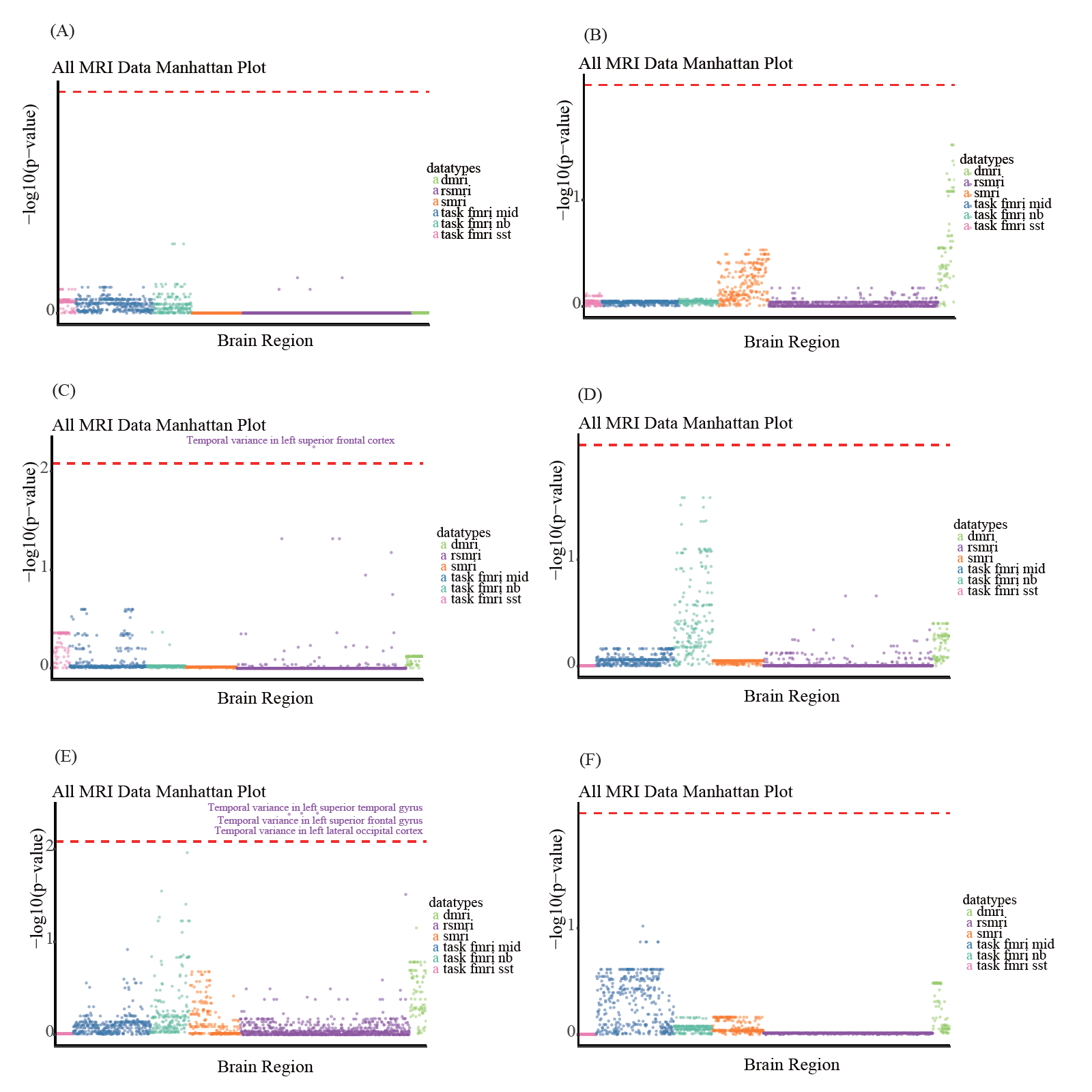

**Fig. S3. The results of linear model at baseline after matching**. (A) the Manhattan plot of the baseline linear model results of ADHD with p(FDR)<0.05/6 (red line); (B) the Manhattan plot of the baseline linear model results of anxiety with p(FDR)<0.05/6 (red line); (C) the Manhattan plot of the baseline linear model results of conduct with p(FDR)<0.05/6 (red line); (D) the Manhattan plot of the baseline linear model results of depression with p(FDR)<0.05/6 (red line); (E) the Manhattan plot of the baseline linear model results of oppositional defiant with p(FDR)<0.05/6 (red line); (F) the Manhattan plot of the baseline linear model results of somatic with p(FDR)<0.05/6 (red line).

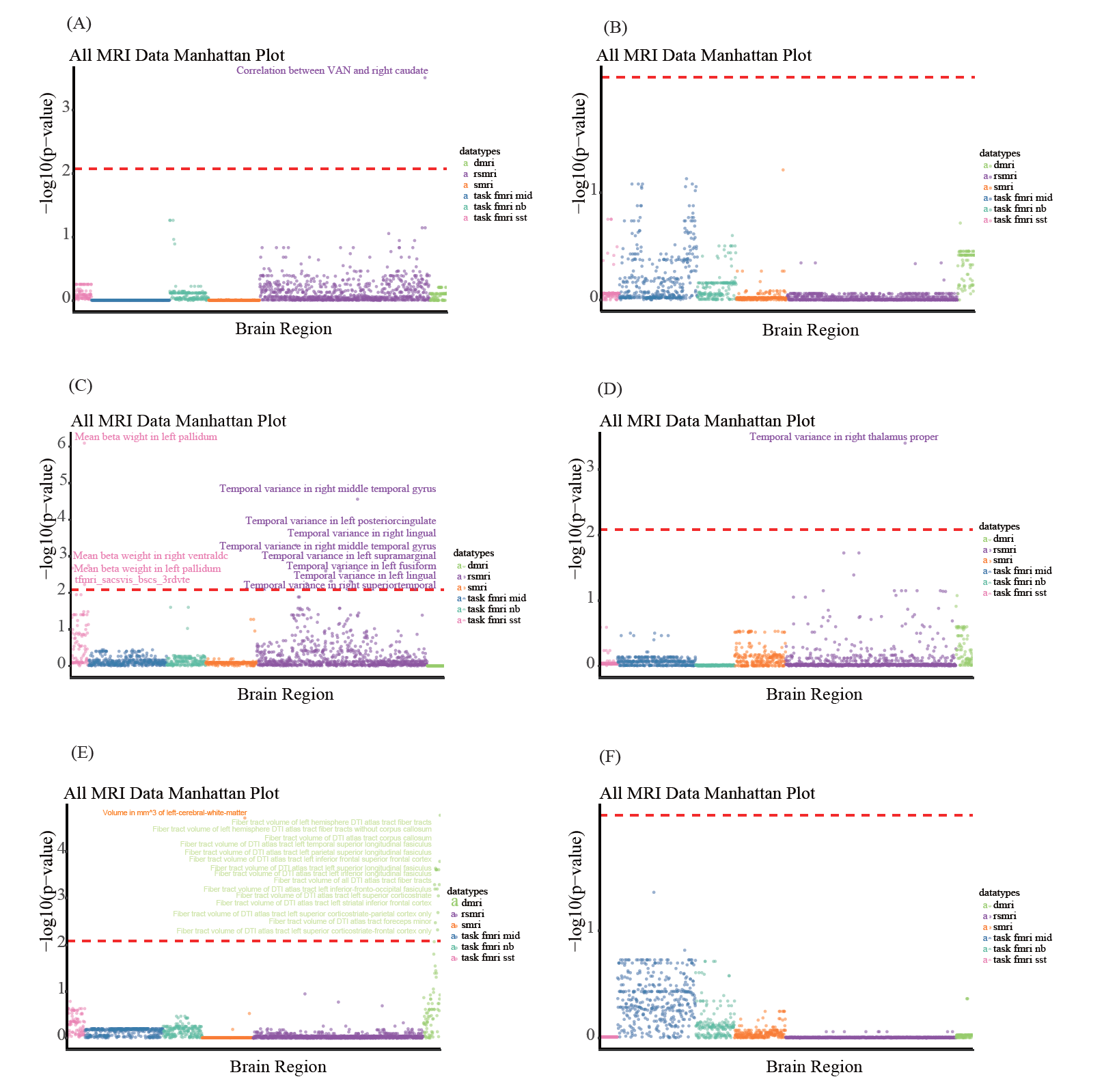

**Fig. S4. The results of linear model of changes after matching**. (A) the Manhattan plot of the change model results of ADHD with p(FDR)<0.05/6 (red line); (B) the Manhattan plot of the change model results of anxiety with p(FDR)<0.05/6 (red line); (C) the Manhattan plot of the change model results of conduct with p(FDR)<0.05/6 (red line); (D) the Manhattan plot of the change model results of depression with p(FDR)<0.05/6 (red line); (E) the Manhattan plot of the change model results of oppositional defiant with p(FDR)<0.05/6 (red line); (F) the Manhattan plot of the change model results of somatic with p(FDR)<0.05/6 (red line).

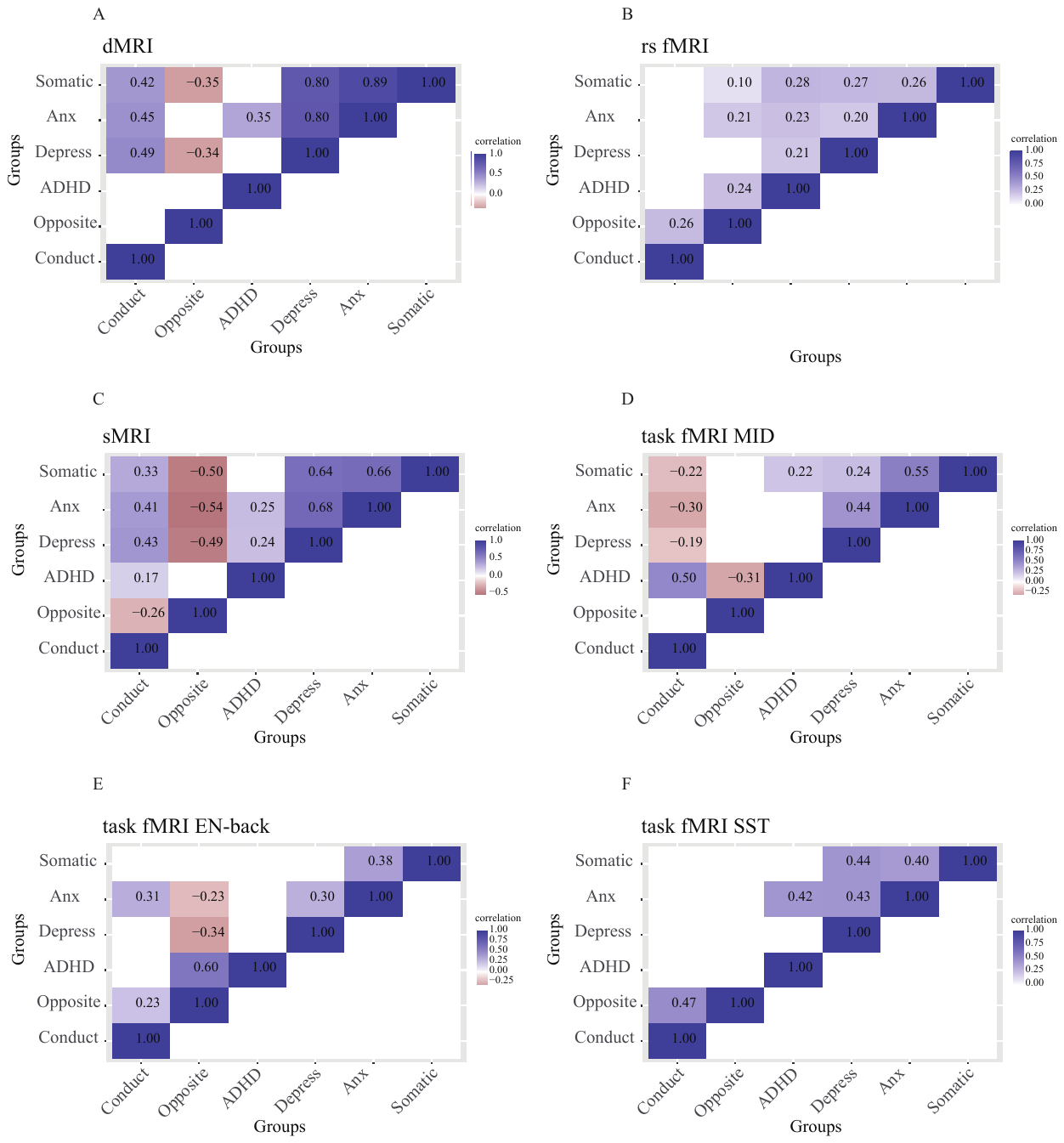

**Fig. S5. The correlation matrices between mental health problems in different modalities at baseline after matching** (blue is positive and red is negative). Only significant correlations between t-values from the baseline linear models at p(FDR)<0.05/6 are shown. (A) DTI; (B) rsfMRI; (C) sMRI; (D) MID task fMRI; (E) EN-back task fMRI; (F) SST task fMRI.

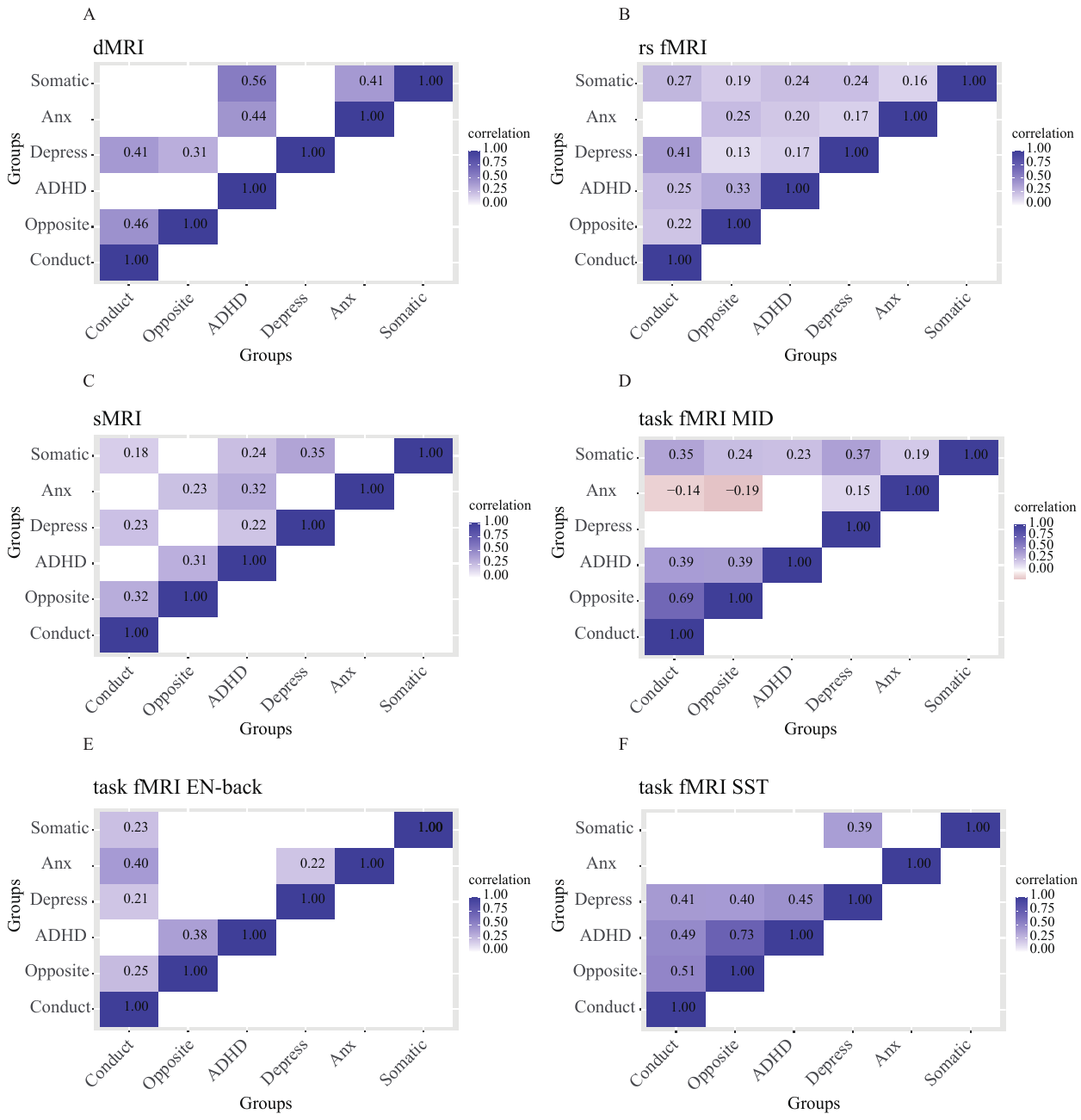

**Fig. S6. The correlation matrices between mental health problems in different modalities of changes after matching** (blue is positive and red is negative). Only significant correlations between t-values from the change linear models at p(FDR)<0.05/6 are shown. (A) DTI; (B) rsfMRI; (C) sMRI; (D) MID task fMRI; (E) EN-back task fMRI; (F) SST task fMRI.

**Table S1. The MRI measures we selected.**

| Modalities | Number | Measures |
| --- | --- | --- |
| sMRI | 258 | Volume of subcortical ROIs |
|  |  | Cortical thickness and cortical area of cortical ROIs |
| rsfMRI | 856 | Correlation within and between cortical networks |
|  |  | Correlation between cortical networks and subcortical ROIs |
|  |  | Temporal variance in subcortical ROIs, cortical ROIs and in Gordon parcels |
| dMRI | 84 | Fractional anisotropy (FA) within DTI atlas tract |
|  |  | Fiber tract volume within DTI atlas tract |
| SST | 90 | Mean beta weight for SST correct stop vs correct go, incorrect stop vs correct go, correct stop vs incorrect go and incorrect stop vs incorrect go in subcortical and cortical ROIs |
| MID | 392 | Beta weight for MID all anticipation of reward/loss vs neutral contrast in subcortical and cortical ROIs |
|  |  | Beta weight for MID all reward/loss positive vs negative feedback contrast in subcortical and cortical ROIs |
| N-back | 196 | Mean beta weight for n-back 2 back vs 0 back contrast in subcortical and cortical ROIs |
|  |  | Mean beta weight for n-back face vs place contrast in subcortical and cortical ROIs |

**Table S2. The significant features from linear models of conduct problems, depression and oppositional defiant problems at baseline (p< 0.05/6).**

| p_fdr | t_values | Regions | Data_type |
| --- | --- | --- | --- |
| Conduct | | | |
| 0.001824134 | 4.75153583 | Temporal variance in left hemisphere cortical Gordon parcel 148 (L_superiorfrontal) | rs fmri  at baseline |
| Opposite | | | |
| 0.004132195 | 4.513288745 | Temporal variance in left hemisphere cortical Gordon parcel 68 (L_superiortemporal) | rs fmri at baseline |
| 0.004132195 | 4.433702417 | Temporal variance in left hemisphere cortical Gordon parcel 148 (L_superiorfrontal) | rs fmri at baseline |
| 0.004236282 | 4.339346044 | Temporal variance in left hemisphere cortical Gordon parcel 5 (L_lateraloccipital) | rs fmri at baseline |

**Table S3. The significant features from linear models of ADHD, conduct problems, depression and oppositional defiant problems during development (p< 0.05/6).**

| p_fdr | t_values | Regions | Data_type |
| --- | --- | --- | --- |
| ADHD | | | |
| 0.000223236 | 5.163610988 | Average correlation between ventral attention network and ASEG ROI right-caudate | rs fmri during development |
| Conduct | | | |
| 3.83E-07 | -5.894590897 | Mean beta weight for SST correct stop versus incorrect stop contrast in ASEG ROI left-pallidum | task fmri sst during development |
| 1.02E-05 | 5.720304765 | Temporal variance in right hemisphere cortical Gordon parcel 264 (right middle temporal gyrus) | rs fmri during development |
| 0.000406067 | 4.914058101 | Temporal variance in APARC ROI left-posteriorcingulate | rs fmri during development |
| 0.00137408 | -4.176980105 | Mean beta weight for SST correct stop versus incorrect stop contrast in ASEG ROI right-ventraldc | task fmri sst during development |
| 0.001525705 | 4.499135124 | Temporal variance in right hemisphere cortical Gordon parcel 177 (R_lingual) | rs fmri during development |
| 0.001525705 | 4.523364805 | Temporal variance in right hemisphere cortical Gordon parcel 267 (right middle temporal gyrus) | rs fmri during development |
| 0.001634272 | -4.04240753 | Mean beta weight for SST correct stop versus correct go contrast in ASEG ROI left-pallidum | task fmri sst during development |
| 0.002445632 | 4.347898402 | Temporal variance in left hemisphere cortical Gordon parcel 104 (L_supramarginal) | rs fmri during development |
| 0.002996513 | 4.261938953 | Temporal variance in left hemisphere cortical Gordon parcel 132 (L_fusiform) | rs fmri during development |
| 0.003492561 | 4.161676296 | Temporal variance in left hemisphere cortical Gordon parcel 8 (L_lingual) | rs fmri during development |
| 0.003492561 | 4.184100184 | Temporal variance in right hemisphere cortical Gordon parcel 228 (R_superiortemporal) | rs fmri during development |
| Depression | | | |
| 0.00040549 | 5.049359547 | Temporal variance in ASEG ROI right-thalamus-proper | rs fmri during development |
| Opposite | | | |
| 1.67E-05 | 5.215275666 | Fiber tract volume in mm^3 of left hemisphere DTI atlas tract fiber tracts | dmri during development |
| 1.91E-05 | 5.395733569 | Volume in mm^3 of ASEG ROI left-cerebral-white-matter | smri during development |
| 0.00016279 | 4.628048836 | Fiber tract volume in mm^3 of left hemisphere DTI atlas tract fiber tracts without corpus callosum | dmri during development |
| 0.000228331 | 4.470236442 | Fiber tract volume in mm^3 of DTI atlas tract corpus callosum | dmri during development |
| 0.000244733 | 4.303051304 | Fiber tract volume in mm^3 of DTI atlas tract left temporal superior longitudinal fasiculus | dmri during development |
| 0.000244733 | 4.373197625 | Fiber tract volume in mm^3 of DTI atlas tract left parietal superior longitudinal fasiculus | dmri during development |
| 0.000244733 | 4.312958539 | Fiber tract volume in mm^3 of DTI atlas tract left inferior frontal superior frontal cortex | dmri during development |
| 0.000245938 | 4.267434195 | Fiber tract volume in mm^3 of DTI atlas tract left superior longitudinal fasiculus | dmri during development |
| 0.000511991 | 4.047968673 | Fiber tract volume in mm^3 of DTI atlas tract left inferior longitudinal fasiculus | dmri during development |
| 0.000511991 | 4.040749239 | Fiber tract volume in mm^3 of all DTI atlas tract fiber tracts | dmri during development |
| 0.000636166 | 3.941045104 | Fiber tract volume in mm^3 of DTI atlas tract left inferior-fronto-occipital fasiculus | dmri during development |
| 0.000636166 | 3.953433697 | Fiber tract volume in mm^3 of DTI atlas tract left superior corticostriate | dmri during development |
| 0.000891349 | 3.83745166 | Fiber tract volume in mm^3 of DTI atlas tract left striatal inferior frontal cortex | dmri during development |
| 0.002033904 | 3.607772775 | Fiber tract volume in mm^3 of DTI atlas tract left superior corticostriate-parietal cortex only | dmri during development |
| 0.003352477 | 3.455258209 | Fiber tract volume in mm^3 of DTI atlas tract foreceps minor | dmri during development |
| 0.004779966 | 3.33877935 | Fiber tract volume in mm^3 of DTI atlas tract left superior corticostriate-frontal cortex only | dmri during development |

**Table S4. The significant features from linear models of somatic problems during development (p< 0.05).**

| p_fdr | t_values | | Regions | Data_type | |
| --- | --- | --- | --- | --- | --- |
| Somatic | | | | | |
| 0.04172532 | -3.881246495 | Beta weight for MID all anticipation of reward versus neutral contrast in APARC ROI rh-superiortempora | | | task fmri mid during development |

**Table S5.** Demographic data of seven group adolescents at baseline after matching.

|  | ADHD | | | Anxiety | | | Conduct | | |
| --- | --- | --- | --- | --- | --- | --- | --- | --- | --- |
|  | Control | Patients | P value_FDR | Control | Patients | P value_FDR | Control | Patients | P value_FDR |
| Sample size | 2128 | 75 |  | 2500 | 114 |  | 2241 | 52 |  |
| Age (mean ± SD)^1^ | 9.93 ± 0.62 | 10.01 ± 0.62 | 0.35 | 9.93 ± 0.62 | 9.83 ± 0.61 | 0.32 | 9.93 ± 0.62 | 9.81 ± 0.62 | 0.20 |
| Gender Male (%)^2^ | 49.15% | 61.33% | 0.11 | 48.36% | 50.88% | 0.67 | 47.97% | 48.08% | 1.0 |
| IQ (mean ± SD)^1^ | 99.05 ± 13.07 | 95.64 ± 14.86 | 0.11 | 103.62 ± 17.10 | 102.07 ± 17.83 | 0.48 | 102.46 ± 16.55 | 99.33 ± 14.54 | 0.20 |
| EA (mean ± SD)^1^ | 16.81 ± 2.61 | 16.83 ± 2.21 | 0.96 | 17.01 ± 2.56 | 16.69 ± 2.72 | 0.44 | 16.66 ± 2.45 | 15.88 ± 2.76 | 0.20 |

|  | Depression | | | Oppositional | | | Somatic | | |
| --- | --- | --- | --- | --- | --- | --- | --- | --- | --- |
|  | Control | Patients | P value_FDR | Control | Patients | P value_FDR | Control | Patients | P value_FDR |
| Sample size | 2349 | 129 |  | 2500 | 58 |  | 2367 | 173 |  |
| Age (mean ± SD)^1^ | 9.91 ± 0.60 | 9.81 ± 0.61 | 0.08 | 9.93 ± 0.62 | 9.82 ± 0.65 | 0.40 | 9.93 ± 0.62 | 9.87 ± 0.62 | 0.18 |
| Gender Male (%)^2^ | 47.21% | 37.98% | 0.08 | 48.36% | 62.07% | 0.21 | 48.12% | 53.76% | 0.18 |
| IQ (mean ± SD) | 103.36 ± 16.99 | 102.12 ± 16.15 | 0.39 | 103.62 ± 17.10 | 101.53 ± 15.88 | 0.43 | 102.19 ± 15.60 | 100.30 ± 13.54 | 0.16 |
| EA (mean ± SD) | 16.92 ± 2.53 | 16.43 ± 2.86 | 0.08 | 17.01 ± 2.56 | 17.10 ± 2.48 | 0.79 | 17.01 ± 2.56 | 16.86 ± 2.51 | 0.16 |

^1^ t-test

^2^ χ ^2^
